# “Health literacy assessment of primary care patients in Low and Middle Income Countries”

**DOI:** 10.1101/630533

**Authors:** F. Pasha, D. Dreshaj, A. Ismaili, I. Sopjani, J. Brooke, Sh. Dreshaj

## Abstract

**Aim:** To explore health literacy levels of primary care patients, and associations with demographic variables, frequency of visits, hospitalization rates, and self-perception of health.

**Background:** Health literacy is the ability to obtain, read, understand and apply healthcare information to inform decision-making to commence or adhere to treatment. The benefits of a population proficient in health literacy include enhanced communication, adherence to treatment, engagement in self-care, and ultimately improved health with financial savings for healthcare providers.

**Design:** Cross-sectional epidemiological study, reported using STROBE guidelines.

**Materials and Methods:** Data were collected from patients attending a primary care center in Prishtina, Kosovo from August to September 2018. Data collection included the Short Test of Functional Health Literacy in Adults (S-TOFHLA), gender, age, socioeconomic status, education, self-perception of health, number of visits to the doctor and hospital. Data were analyzed with SPSS software (version 20).

**Results:** Participants (n=557) ages ranged from 15 to >65 (mean 27.82) years, were female (57.85%), Albanian (97.13%), with a response rate of 92.83%, 79% were health literate, 9% had moderate health literacy, 12% were health illiterate. Four variables determined health literacy, level of education (p < .01), gender (p = .033), hospitalization rates (p < .05), socioeconomic status of unemployed compared to being a student (p<.01).

**Conclusions:** There remains a need to address health literacy levels in Kosovo, through the development or adaptation of health literacy tools appropriate for this population, which will support and positively impact on patient’s wellbeing. Nurses are the best-placed professionals to implement these tools and support patients with low health literacy.

**Relevance to clinical practice:** Nurses have a key role in implementing health literacy tools and supporting patients by adapting their communication styles in accordance with each patient’s level of health literacy, which will support adherence to advice, safety and treatment outcomes.

**What does this paper contribute to the wider global clinical community?:**

- Health literacy is associated with level of education, gender and socioeconomic status and hospitalization rates of primary care patients.
- A focus on health literacy is essential to address the inequalities of health for those with marginal or inadequate health literacy.
- Nurses are the best-placed health care professionals to support individuals with low levels of health literacy through interventions, and adapting their communication styles.

## INTRODUCTION

Over two decades ago, Nutbeam (2000) identified the importance of health literacy as a relatively new concept in the developing field of health promotion, alongside Kickbusch (2001) who identified the need to address the disparity of the health and education divide. The Institute of Medicine (IOM, 2004) defined health literacy as “the degree to which individuals have the capacity to obtain process and understand basic health information and services to make appropriate health decisions.” The World Health Organization (WHO, 2019) further defined health literacy as “the cognitive and social skills which determine the motivation and ability of individuals to gain access to, understand and use information in ways which promote and maintain good health.” The WHO (2019) definition of health literacy, identifies the need for individuals to acquire a complex set of skills, including reading, listening, understanding and interpretation of health literature to enable them to engage fully in communication with healthcare providers and genuine involvement in informed decision making (Berkman et al., 2011).

Health literacy, however, can be measured by a screening tool (Roy et al., 2019), such as the Functional Health Literacy in Adults (TOFHLA) (Parker et al., 1995), which has been amended to the Short Functional Health Literacy in Adults (S-TOFHLA) (Baker et al., 1999). The S-TOFHLA composes of 4 items assessing numeracy and 36 items assessing reading comprehension, which enables the classification of health literacy of a person or population as inadequate, marginal, or adequate (Baker et al., 1999). The importance of health literacy has recently been acknowledged, and the S-TOFHLA has been widely implemented in various settings and health populations including patients with diabetes (da Rocha et al., 2019), hemodialysis (Brice et al., 2019), cardiovascular disease (Elbashir et al., 2019), infectious diseases (Castro-Sanchez et al., 2016), and carers of older people (da Almeida et al., 2019).

Marginal and inadequate levels of health literacy have been associated with lower engagement in health prevention activities such as adoption of immunization regimes (Castro-Sanchez et al., 2016), and screening for breast cancer by attending a mammography (Fernandez, Larson & Zikmund-Fisher, 2019). Poor health literacy has also been associated with lower engagement in self-management behaviours such as medication adherence in patients with type 2 diabetes (da Rocha et al., 2019), and adherence and poorer inhaler technique in patients with chronic obstructive pulmonary disease (O’Conor et al., 2019). Low health literacy has also been associated with negative health outcomes such as mortality in older adults (Bostock & Steptoe, 2012), and increased use of healthcare services (Bains & Egede, 2011) and hospitalization for people with long term conditions, such as heart failure (Wu et al., 2013) and cancer (Cartwright et al., 2017).

Health literacy is a global public health concern, as low health literacy has consistently been associated with people of older age, low education levels, low income and a lack of access to the internet (Protheroe et al., 2017), and follows a social gradient, which reinforces socioeconomic health inequalities (Harris et al., 2015; Kickbusch et al., 2013). Low health literacy has an economic cost due to the increased need for healthcare provision; in America and the United Kingdom, the estimated cost ranges between 3-5% of the annual health budget (Eichler, Wieser & Brugger, 2009; Lamb & Berry, 2014). The World Health Organization acknowledged the need to address the issue of equitable health literacy with an explicit call for action in the Seventh Global Conference on Health Promotion in 2009. Since this date guidelines, policies, scientific statements and practical tools address the social disparity and low levels of health literacy.

The World Health Organization Europe in the Health 2020 policy framework (WHO Europe, 2013), identify the first priority of investing in health promotion across the life span and empowering individuals through a strong focus on improving health literacy. Scientific statements support this approach, such as the American Heart Association scientific statement to address the adverse cardiovascular outcomes associated with low levels of health literacy (Magnani et al., 2018). Practical tools to support health literacy have included the development of health literacy tool kits, these have been developed by governing bodies of health education, such as NHS Education for Scotland and Health Education England with the support of CHL Foundation. Beyond Europe, the World Health Organization has produced a health literacy tool kit for low and middle-income countries, which provides information sheets to empower communities and strengthen health systems (Dodson, Good & Osbourne, 2015).

Health literacy in low and middle-income countries is limited, focus on health literacy and interventions to improve health literacy have focused on high-income countries, this needs to be addressed (Dodson, Good & Osbourne, 2015). Kosovo, is a low and middle-income country, with one study identifying limited health literacy in 43.8% of a cross-sectional sample of primary care users (Toci et al., 2014) this is lower compared to the average across eight European countries of 47% (Sørensen et al., 2015). The work by Toci et al. included the exploration of associations with health literacy including sociodemographic and economic status, self-perceived health, and chronic morbidity (Toci et al., 2013; Toci et al., 2015). The aim of this study is further this initial work to explore the association between health literacy levels, education, socioeconomic status, presence of chronic diseases, self-perception of health, frequency of visits to the doctor and hospitalization rates of primary care patients.

## METHODS

### Setting

The setting for this study was seven primary care centres in Prishtina, Kosovo. Kosovo, a country in southeast Europe in the Western Balkans, has a population of 1.7 million, and the capital Prishtina has 207,313 inhabitants. The provision of primary healthcare in Prishtina is through family medical centres, which cover both the urban and rural areas of the capital city. During 2017, family medical centres of Prishtina provided over one million medical consultions and over two and a half million other medical services. Primary healthcare in Kosovo provides initial diagnoses and patient care for 80-90 % of all cases, serving as gatekeepers for secondary and tertiary healthcare. Family medical centres are also responsible for diagnosis and treatment, including minor surgery, drug management, immunization, emergency care and stabilization of emergency patients, maternal and child healthcare, reproductive health services, including antenatal and post-natal care, as well as family planning and treatment of sexually transmitted diseases.

### Participants

Consecutive case sampling was applied; every third consecutive patient who attended a GP appointed was approached to participate in the study. The inclusion criteria was broad, and included all patients 15 years and older, who spoke or understood English. The exclusion criteria were emergency cases, or patients with mental or physical health disabilities that could not complete the required information, and those who refused.

### Variables

Variables collected in this study included: demographic data (gender and age); level of education (unschooled, or completion of primary school, high school, university, or masters/doctoral studies); socioeconomic status (employed, unemployed, student, retired and social aid); documentation of co-morbidities; self-perception of health (healthy, moderately healthy, sick); frequency of visits to the doctor; hospitalization rates of primary care patients, and health literacy of patients.

### Health literacy

Health literacy was measured by the Short Test of Functional Health Literacy in Adults [S-TOFHLA], which consists of 2 prose passages with 36 items, with a Cronbach’s alpha of 0.97 (Baker et al., 1999). The S-TOFHLA is a practical measure of functional health literacy, with the classification of individual’s function health literacy as inadequate, marginal or adequate (refer to Table 1).

**Table 1:** S-TOFHLA Functional Health Literacy Level

| Level | S-TOFHLA Score | Functional health literacy description |
| --- | --- | --- |
| Inadequate functional health literacy | 0-16 | Unable to read or interpret health texts. |
| Marginal functional health literacy | 17-22 | Has difficulty reading and interpreting health texts. |
| Adequate functional health literacy | 23-36 | Can interpret and read most health texts. |

### Data collection

Data collection occurred at primary care centres from August to September 2018. The collection of data occurred by randomly selected general practitioners (GPs). All GPs received detailed instructions and training on recruitment of participants, how to implement the scripted introduction, test administration, and evaluations. GPs were informed of the administration criteria of a 7-minute time limit for the completion of the S-TOFHLA, this was adhered to by using the stopwatch on a smartphone. In total 30 GPs were recruited to complete data collection, each GP was provided with a target of recruiting 20 participants over 20 working days to capture a cross-sectional sample of a distinct period of time.

The S-TOFHLA has been validated for both English and Spanish speaking populations (Aguirre et al., 2005). Therefore, our sample population included only patients who could speak and understand the English language, thus using S-TOFHLA English version for testing. Following the completion of the S-TOFHLA, GPs documented gender, age, socioeconomic status, self-perception of health education, language, number of visits to the doctor and hospitalization rates, through a fourteen question multiple choice questionnaire on patient’s general information.

### Ethical consideration

The Ethical Committee of the Main Family Medical Center in Municipality of Prishtina, Republic of Kosovo (protocol number 453, approval date 12.02.2018), provided ethical approval and continuous monitoring of this study. Participants received detailed information of the study, prior to the voluntary provision of informed consent. For participants still legally considered as a minor, parental consent was obtained. Confidentiality of participants was maintained during data collection, analysis and reporting of the results.

### Data analysis

Data were analyzed with the IBM Statistical Package for the Social Sciences Statistics version 20.

Gender was the only nominal data, all other data was continuous in nature. Descriptive statistics were used to determine gender, age groups, nationality, and level of education. The S-TOFHLA score was divided into three categories, and participants’ were defined as having adequate health literacy if they scored between 23-36, as marginal health literacy if they scored between 17-22, and inadequate health literacy if they scored is 0-16 [refer to Table 1]. The statistical analysis included Levene’s test for equality of variances to determine the significant differences of health literacy with gender and age, a oneway ANOVA to determine the significant differences of health literacy and social status, and education.

### Rigour

The Republic of Kosovo has a population of 1.7 million inhabitants, while our sample size population included 557 participants. Sample size power calculation in 95% confidence interval and 5% common standard deviation is 1.0 or 100%. The power calculation suggested the need to recruit 384 participants to ensure the power of this study.

## RESULTS

The response rate was 92.83%, 557 participants out of a possible 600, consisting of 57.85% females and 42.15% males, coming from urban and rural areas of Municipality of Prishtina. The highest participant age range was 16-25 and 26-35 years, consisting of 29.69%, and 28.54% respectively. The majority of the participants were Albanians (97.13%) and the remaining participants were Bosnians, Roma-Egyptians and Turkish. From all participants 0.74% were unschooled, 4.44% finished primary school, 29.94% high school, 54.53% had a university degree and 10.35% a post-graduate qualification (refer to Table 2).

**Table 2:**
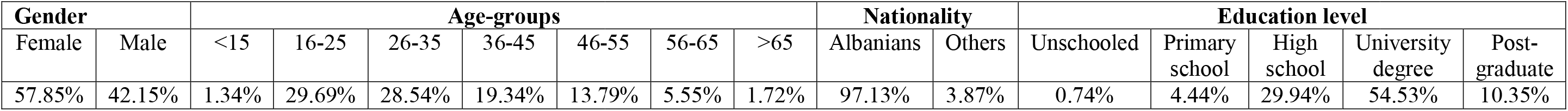
Study population characteristics

The scoring criteria of the S-TOFHLA identified 79% of our population had adequate health literacy level, 12% had inadequate health literacy level, and 9% had marginal health literacy.

**Figure 1:**
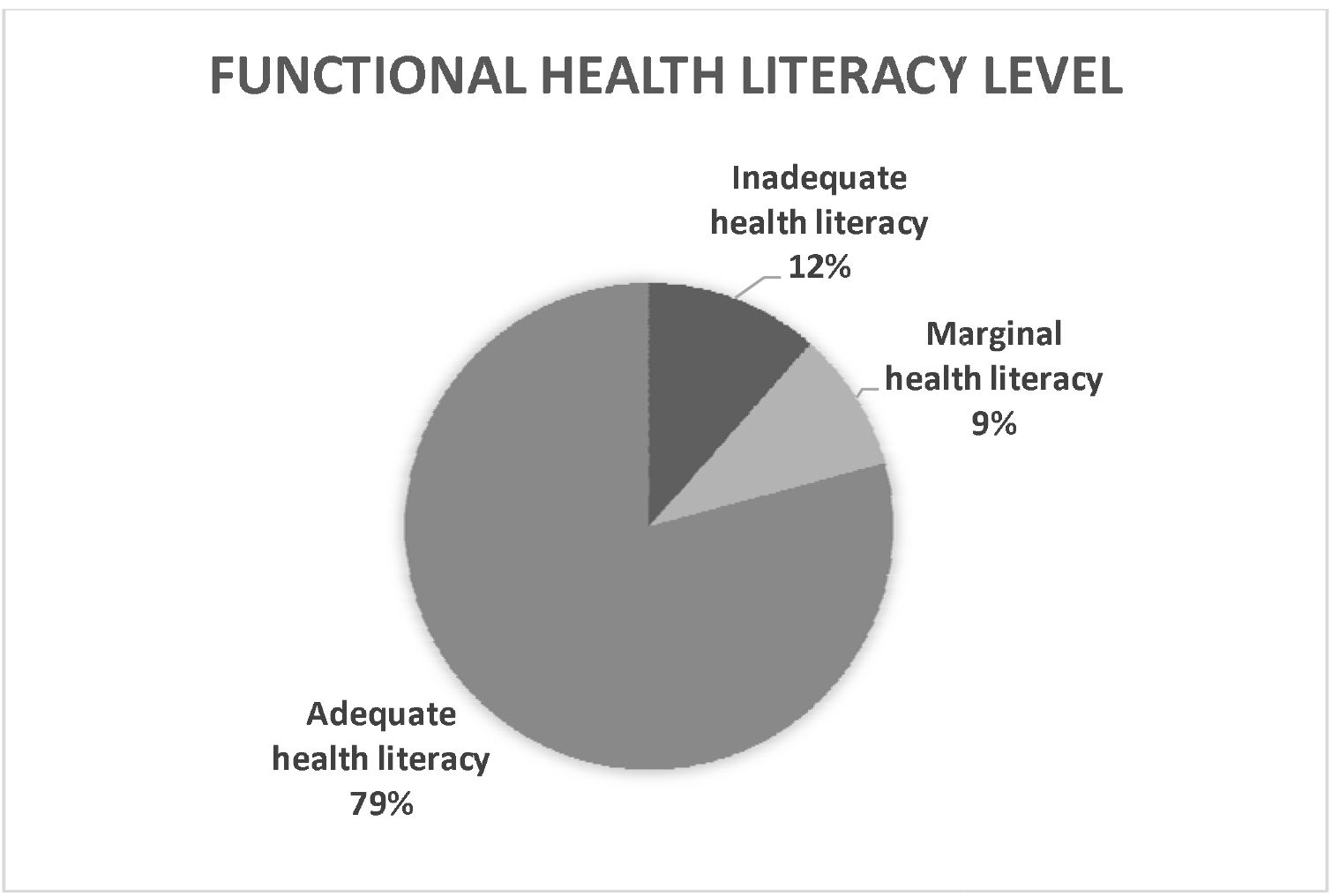
Health literacy test performance

A statistically significant difference (p = .033) between females (M = 28.37, SD = 7.251) and males (M = 26.96, SD = 7.863) and health literacy was identified. There were no significant differences between age groups and health literacy.

The results of a one-way ANOVA demonstrated statistically significant differences (p < .01) in health literacy and being unemployed (M = 26.5, SD = 7.972) or a student (M = 29.19, SD = 7.238), with higher levels of health literacy in the latter group. No other significant differences between socioeconomic status and health literacy were identified (refer to Figure 2).

**Figure 2:**
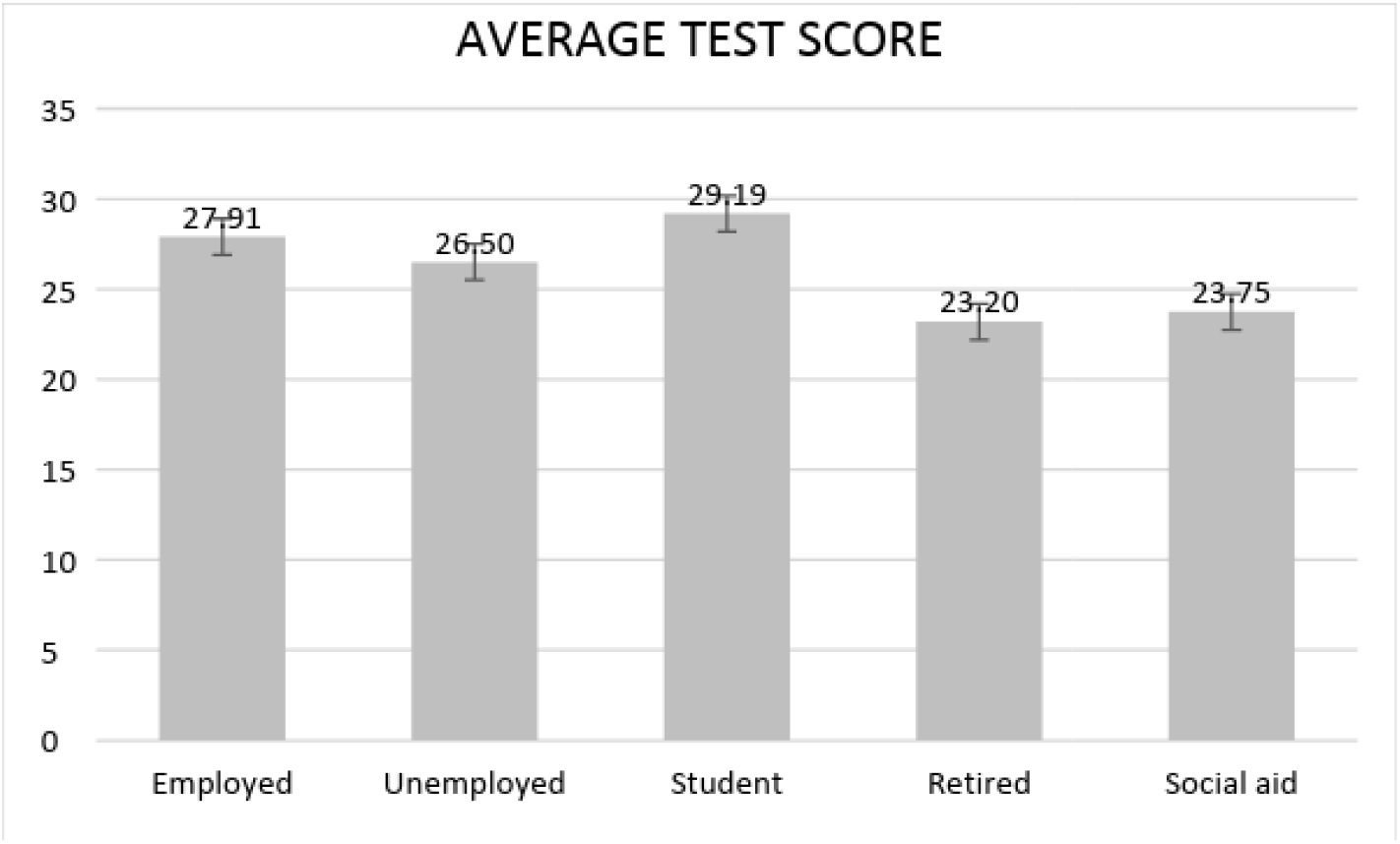
Health literacy and social status

The results of a one-way ANOVA demonstrated statistically significant differences (p < .01) in health literacy and patients with different levels of completed education. Further exploration of results through Tukey’s HSD (honest significant difference) demonstrated the significant difference was between those who were unschooled (M = 18.75, SD = 8.5) or with only a primary school education (M = 24.38, SD = 9.016), who had lower health literacy, than those with a Bachelor Degree (M = 28.69, SD = 7.309) or Masters/Doctoral degree (M = 29.63. SD = 6.571), who had higher health literacy (refer to Figure 3).

**Figure 3:**
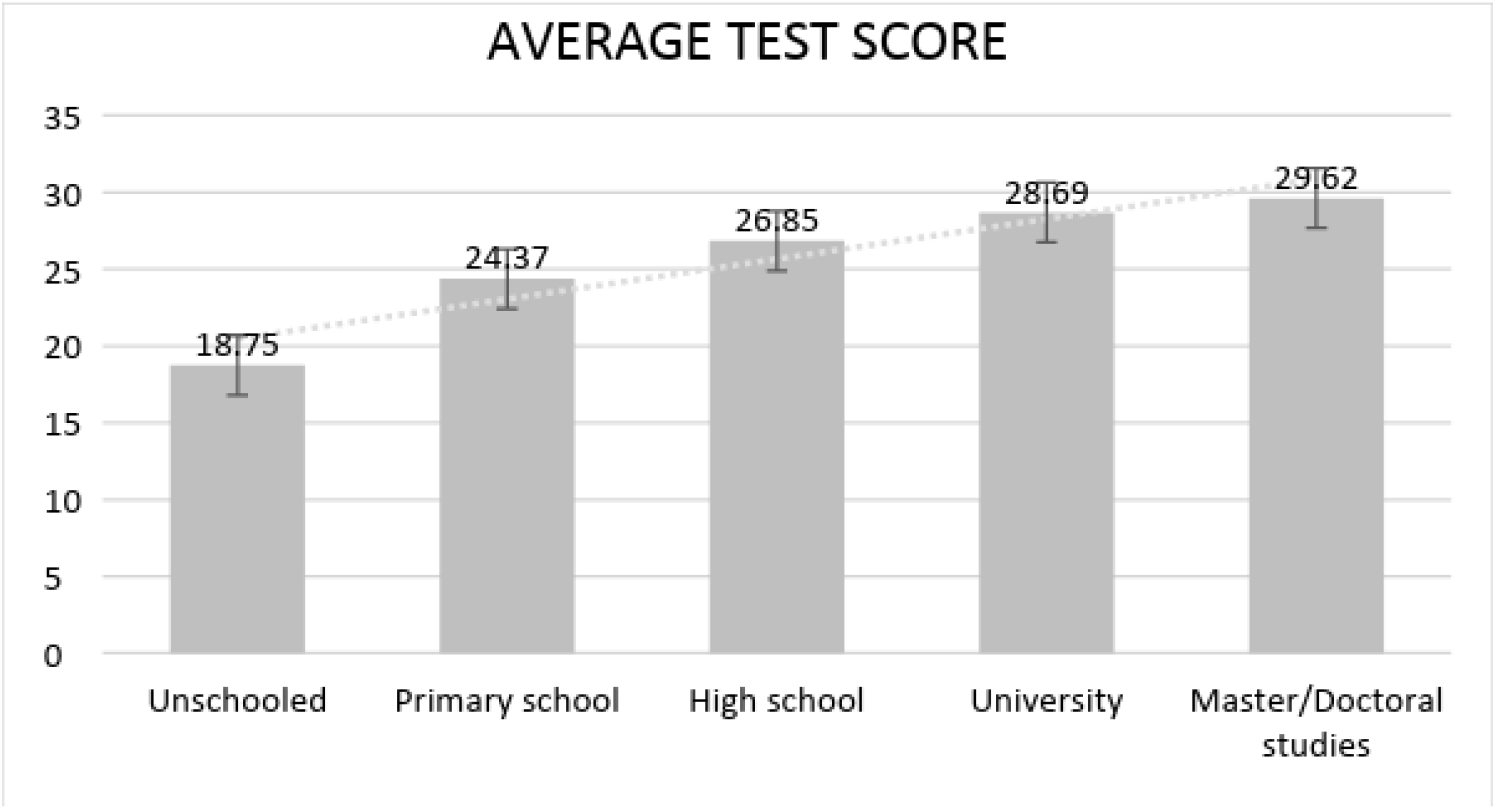
Health literacy and education

While there were no significant differences in health literacy and number of visits to the doctor, there were statistically significant differences (p < .05) between those who have never been hospitalized (M = 29.43, 6.764) and those who have been hospitalized at least once in their life (M = 27.73, SD = 7.739), the latter group had lower health literacy.

Further analysis identified that patients who were never hospitalized were significantly more likely (p < .01) to perceive themselves as healthy people, compared to those who have been hospitalized once, twice or more times. Similarly, those who visited doctor less often were again significantly more likely (p < .01) to report their health status as “healthy”.

Results revealed that there are statistically significant differences (p < .01) in frequency of doctor visits between patients with and without a chronic disease. As expected, those with a chronic disease are significantly more likely to pay more visits to the doctor within a year, when compared to those who do not suffer from a chronic disease.

## DISCUSSION

Our study explored the level of health literacy of primary care patients in the Municipality of Prishtina, Kosovo, with 79% of participants demonstrating an adequate health literacy, 9% marginal health literacy and 12 % inadequate health literacy. These results are not comparable with the previous work of health literacy in Pristine, Gjakove and Prizren, which identified limited health literacy in 43.8% of their sample (Toci et al., 2014). However, the results of Toci et al. (2014) are comparable with a European study across eight countries, which identified 47% of individuals with limited, insufficient or problematic health literacy measured by the Health Literacy Survey – European Union questionnaire (HLS-EU-Q) (Sørensen et al., 2013; 2015). Although, inadequate health literacy can differ substantially across countries, such as 1.6% in the Netherlands to 26.9% in Bulgaria (Sørensen et al., 2015). These substantial differences are replicated in Low and Middle Income Countries (LMICs), across 14 countries in Sub-Saharan Africa health literacy ranged from 13% in Ethiopia to 64% in Namibia (McClintock et al., 2017).

In our study, we identified significant differences of health literacy among male and female participants, education, socioeconomic status and hospitalization rates. Higher levels of health literacy were identified among female primary care patients than male patients. However, in Sub-Saharan Africa the reverse was found, with higher levels of literacy identified among men (39.2%) compared to women (34·1%) (McClintock et al., 2017). In a longitudinal study over the life course, being male was a significant predictor of health literacy with data collected from 1957-2011 (Clouston, Manganello & Richard, 2017). Our results are comparable to a study completed in Korea, which demonstrates the association between health literacy and gender is complex (Lee, Lee & Kim, 2015). On items such as of how to understand and fill out medical forms and understand directions on medication bottles, males compared to females demonstrated an inadequate understanding (Lee, Lee & Kim, 2015). A further consideration of the different results from these studies may be the result of the role of women in Sub-Saharan Africa, and the opportunities of education post 1960, and the importance of further variables such as education, socioeconomic status and hospitalization rates.

In our study, the level education of primary care patients significantly impacted on their health literacy, with a positive trajectory from unschooled to completion of doctoral level studies. These results are similar to Sub-Saharan countries, as only 9% of adults who completed some primary school education demonstrated adequate health literacy, compared to 69% of adults who completed some secondary school education and 84% who completed all their secondary education (McClintock et al., 2017). The importance of completing secondary school education on health literacy is evident, however for those who have not, the importance of educational interventions to support the development of adequate health literacy is essential. Educational health literacy interventions can enhance health literacy and health-promoting behaviours (Bayati et al., 2018). Nurses and other healthcare professionals need to be competent in supporting and delivering health literacy education from the resources that are available (Loan et al., 2017), such as the World Health Organization has health literacy tool kit for low and middle-income countries (Dodson, Good & Osbourne, 2015).

Another significant variable that predicted health literacy in our study, which was related to education, was economic status as students scored higher compared to those unemployed. Socioeconomic status is recognized to impact and create inequalities in health and those with low socioeconomic status have lower life expectancy and higher morbidity (Mackenbach, 2012; Mackenbach et al., 2008). Therefore, the relationship between health literacy and socioeconomic status is complex, as health literacy has been identified as a possible mediator between the relationship of socioeconomic status and health beliefs and behaviours relating to cancer (Adams et al., 2013). More recently in a systematic review, health literacy has been identified to mediate the relationship between socioeconomic status and health disparities and inequalities (Stormacq, van den Broucke & Wosinski, 2018). This may be exacerbated by health care professional’s recognition of health literacy, as nurses tend to overestimate patient’s health literacy (Dickens et al., 2013). Interventions to increase levels of health literacy knowledge and understanding for nurses to support those with low health literacy is an approach that could create greater equity in access to health care services and the health of individuals (Dickens et al., 2013; Stormacq, van den Broucke & Wosinski, 2018).

Our study identified a significant relationship between lower levels of health literacy and rates of hospitalization. The relationship between lower health literacy and increased hospital admission, and readmission within 30 days has been well documented (Baker et al., 2002; Mitchell et al., 2012; Cimasi, Sharamitaro & Seiler, 2013). Although, a recent study exploring health literacy of adults following a recent hospital admission found no association between inadequate health literacy and increased hospital admissions (Jessup et al., 2017). However, interventions to improve health literacy levels can support patients across the health literacy spectrum, through interventions at nurse-patient level, system-patient level and community-patient level (Sudore & Schillinger, 2009). Nurse-patient level interventions involve adapting communication styles from an understanding of the patient’s level of health literacy, including language, medical terminology and step-by-step information on complex instructions, limiting each interaction to three key points and assessing for comprehension (Adams et al., 2013; Hersh, Salzman & Snyderman, 2015). The investment in understanding and supporting a patient through understanding their level of health literacy will ultimately increase their adherence to complex health regimes and their ability to self-manage their long term conditions improve their outcomes (Marshall, Sahm & McCarthy, 2012). Therefore, a core skill of nursing is to support patients by understanding of their level of health literacy and adapting their communication styles in a person-centered approach (Wittenberg et al., 2019).

### Limitations

The limitation within are study, are firstly this was a single site study in municipality of Prishtina, the capital city of Kosovo, thus we lack information from other regions. Secondly, S-TOFHLA does not assess all aspects of health literacy, such as spoken health literacy, which may impact on external validity of our study.

## CONCLUSION

In our study, three quarters of the primary care patients in Prishtina, Kosovo had adequate health literacy, which was associated with being female, those with higher education attainments, and those who had never had a hospital admission. Therefore, our study identified primary care patients with marginal and inadequate health literacy, which was associated with either being male, unemployed or had an admission to hospital. Therefore, there is a clear need to identify and support primary care patients with marginal and inadequate health literacy, to support and improve their health literacy, and deliver health advice at a level they can understand to improve their health and health outcomes, and reduce pressure of primary and secondary/tertiary healthcare provision.

## RELEVANCE TO CLINICAL PRACTICE

Nurses are the best-placed healthcare professionals to challenge the health inequalities of patients with marginal and inadequate health literacy. All nurses need to understand health literacy and the impact of inadequate and marginal health literacy on the health of their patients. Therefore, there is a need for nurses to be competent in supporting and delivering interventions to improve levels of health literacy. Nurses also need to be able and competent to assess patients health literacy and adapt their communication styles to support patients with low health literacy to empower them to adopt the necessary behaviours to support and improve their health.

